# Rapid and repeated evolution of the pigmentation patterns in reef fishes

**DOI:** 10.1101/2025.02.21.639431

**Authors:** Bruno Frédérich, Laurent Mittelheiser, Amandine Gillet, Jennifer R. Hodge, Vincent Laudet, Alex Dornburg

## Abstract

Pigmentation patterns are integral to animal biology^1–3^ and uncovering the mechanisms driving their diversification is essential for determining the evolutionary principles that shape this fundamental aspect of biodiversity^4–7^. Coral reef fishes are particularly notable for their extraordinary pattern diversity, ranging from simple spots and stripes to intricate, maze-like designs. Despite over a century of investigation, the evolutionary processes that govern the diversification of these pigmentation patterns remain one of the most persistent unresolved questions in evolutionary biology. Here, we investigate the relationship between pattern diversity, species richness, and geography across six iconic families of pattern-diverse coral reef fishes. Utilizing time-calibrated phylogenies, we reveal constant disparity of pigmentation patterns across globally variable reef fish communities^8^. We find strong evidence for a positive correlation between pattern diversity and species richness, with a high divergence of pigmentation patterns in sympatry that highlights the role of these patterns in speciation and phenotypic differentiation. Moreover, our findings support the stages model of adaptive radiation^9^, revealing that most pigmentation pattern diversity has emerged in evolutionary history. These results demonstrate that the evolutionary history of pigmentation patterns in reef fishes is characterized by a combination of rapid and constrained phenotypic diversification that has likely played a crucial role in their speciation dynamics.

---

Color patterns are fundamental to the survival of organisms across the Tree of Life, often forming the basis for camouflage, mimicry, predator deterrence, and communication within and between species^1^. The diversity of animal color patterns also plays an important role in the formation of reproductive barriers, facilitating speciation in lineages as disparate as butterflies and cichlids^10,11^. The teleost fishes that comprise tropical coral reef fish communities exemplify pigmentation pattern diversity, exhibiting some of the most striking and diverse color patterns of all living vertebrates^5^. However, the evolutionary processes that govern the diversification of these patterns are among the most persistent, unresolved questions in evolutionary biology. This knowledge gap limits our understanding of the relationship between pigmentation patterns, ecological adaptation, and speciation. Resolution of this question hinges on whether the diversification of pigmentation patterns is primarily driven by local ecological pressures, leading to marked dissimilarity between ecoregions, or if it is more strongly influenced by speciation processes, leading to rapid diversification among closely related taxa and, consequently, similarities in color pattern diversity across biogeographic regions with heterogeneous species assemblages.

Tropical coral reefs across biogeographic regions, such as the Caribbean and Indo-Pacific, differ significantly in key characteristics, including trophic structure, faunal composition, and species richness^12^. Accordingly, fish clades within these regions may also exhibit notable pigmentation pattern dissimilarities, attributed to ecological or sexual selective pressures. However, the pace at which fish pigmentation patterns have diversified in response to these forces remains an open question. Regional differences in fish assemblages largely reflect historical biogeography^8,13^. Consequently, varying environmental conditions and competitive interactions may have driven divergence in pigmentation pattern over time, leading to a gradual accumulation of pigmentation pattern diversity within fish assemblages. Additionally, adaptive radiation theory predicts traits related to species recognition and communication are the last to diversify in radiations, following initial divergence along habitat axes and morphological specialization related to trophic resources^9^. With strong evidence for extensive speciation in reef fishes across most tropical oceans during the Pliocene^14–18^, this scenario suggests that pigmentation pattern diversity would disproportionately evolve near the tips of phylogenies, driven by the reinforcement of reproductive barriers between closely related species. These competing hypotheses echo the classic “cradles” versus “museums” debate in evolutionary biology^19^ and represent distinct views on the tempo and mode of pigmentation pattern diversification. Determining which of these perspectives holds more explanatory power is crucial for understanding the evolutionary dynamics shaping one of the most iconic traits in coral reef ecosystems.

Here we investigate the key factors shaping pigmentation pattern diversity in coral reef fishes by combining time-calibrated molecular phylogenies, biogeographic data, and comprehensive analyses of pigmentation patterns in six ecologically diverse and globally representative fish families: surgeonfishes (Acanthuridae), butterflyfishes (Chaetodontidae), snappers (Lutjanidae), goatfishes (Mullidae), angelfishes (Pomacanthidae), and damselfishes (Pomacentridae). We first test whether species richness explains pigmentation pattern diversity across biogeographic regions. Next, we investigate how pigmentation pattern disparity varies among reef fish clades across different global regions, providing an assessment of the degree to which ecological variance between regions explains standing patterns of trait disparity. Finally, we evaluate expectations from the theoretical stages of adaptive radiation to determine its relevance in explaining the observed patterns of coral reef fish pigment pattern diversity. Collectively, these findings reveal the evolutionary mechanisms driving pigment pattern diversification, providing new insights that resolve critical aspects of this long-standing question in evolutionary biology.

## Pigmentation pattern diversity is correlated with species richness

Divergences in pigmentation patterns often play a critical role in speciation^20,21^, and recent evidence has suggested an association between motif divergence and genetic divergence among some reef fish populations^22,23^. This association raises the possibility that pattern diversity may be correlated with species richness within biogeographic regions. To test this hypothesis, we annotated 30 pigmentation motifs using images of 918 fish species from the six coral reef fish families, ensuring coverage of at least 75% of the species diversity within each family (**Extended Data Table 1**). These motifs included well-established categories such as stripe patterns (horizontal, diagonal, vertical, labyrinthine), spot patterns (spot and eyespot), and others (saddle-like, blotch, monotone). The head, trunk, and tail regions of each fish were binary coded to indicate the presence of each motif, distinguishing between single occurrences and repeated patterns. We then summarized the data using principal coordinates analysis (PCoA) for each family. Quality assessments confirmed that eight to ten principal coordinate axes were sufficient to capture the phenotypic diversity of each family without significant loss of information (see Methods for details). Pigmentation data were integrated with geographic information from FishBase^24^ and GBIF^25^, and fish species were categorized into five major ecoregions according to the classification by Kulbicki and colleagues^13^: Atlantic (including the Mediterranean Sea), Western Indian (including the Red Sea), Central Indo-Pacific, Central Pacific and Tropical Eastern Pacific (**Extended Data Table 1**).

We quantified the diversity of pigmentation patterns present in each biogeographic region by measuring the *Richness* of motif diversity, defined by the hypervolume occupied by each family in the pigmentation pattern space^26,27^. We find a strong, significant, and positive relationship between species richness and motif diversity per ecoregion (**Figure 1**). These results suggest that species differentiation is tightly associated with motif diversity, regardless of regional species-richness patterns, highlighting the central role of pigmentation patterns in the speciation of coral reef fishes. This also supports the long-established use of pigmentation patterns and color variation in identifying reef fish species^20^.

**Figure 1.**
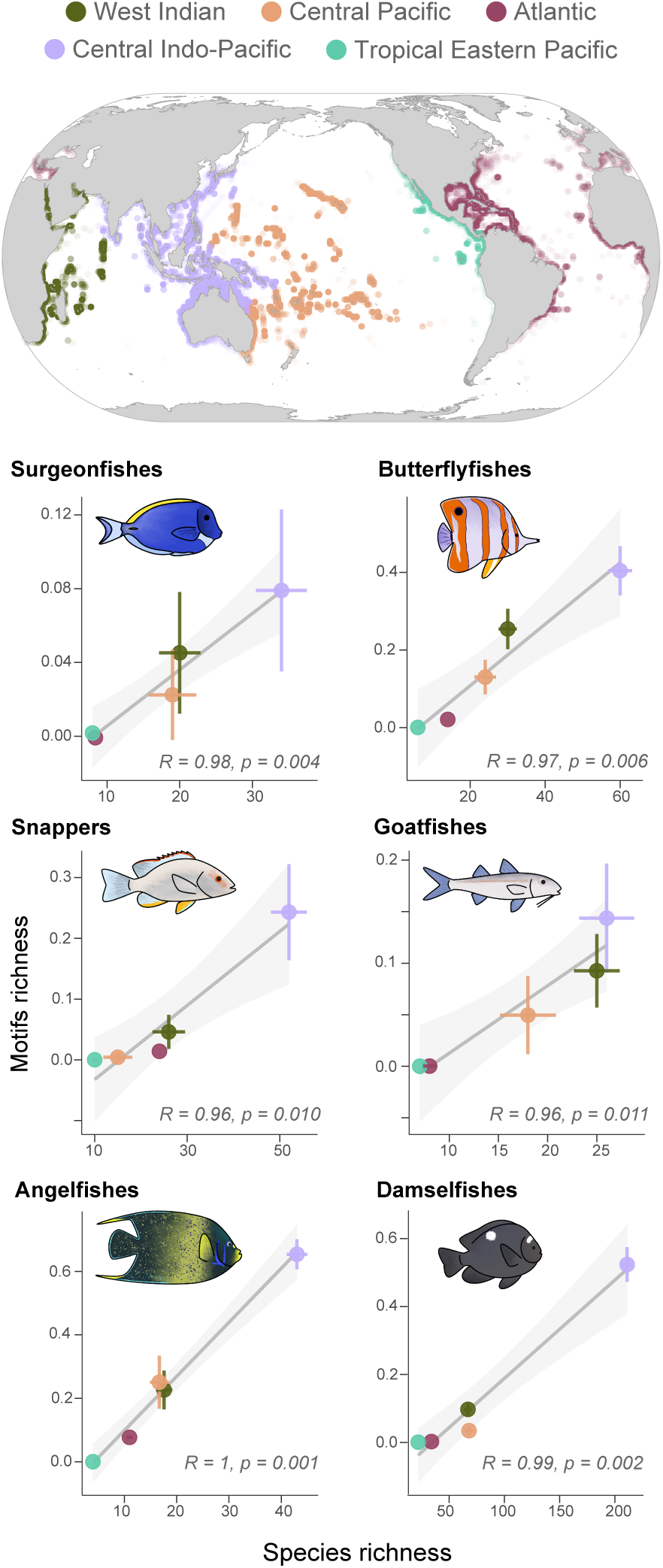
Relationships between motif richness and species richness from the five main ecoregions. The richness of pigmentation patterns (i.e. motif richness) is directly proportional to the number of species present in a region. Tests of correlation are all significant and Pearson’s R coefficients are close to a maximum value of 1.

We also assessed the distribution of species within each hypervolume using two additional metrics: *Divergence* and *Evenness*. *Divergence* measures the distance of species from the center of the group of species forming the occupied space, with high values indicating substantial differentiation from the average pigmentation pattern of the group. *Evenness* reflects the regularity of species distribution across the occupied hypervolume, with higher values indicating a more uniform spread in morphospace^26^. Across all geographical groups and families, both divergence and evenness were generally high, often approaching the maximum value of 1 (**Extended Data Table 2**). The lowest values of *Evenness* are observed in the damselfishes (from 0.63 in the Atlantic Ocean to 0.70 in the Tropical Eastern Pacific and Central Pacific). Similarly, angelfishes had the lowest value of *Divergence* in the Tropical Eastern Pacific (0.65). However, unlike the relationship between motif diversity and species richness, divergence and evenness in pigmentation patterns were generally not correlated with species richness (all correlation tests: *p*>0.1, except a significant negative relationship between divergence and species richness in surgeonfishes (*p*=0.04); see **Extended Data Table 3**). Similarities in geographic patterning of divergence and evenness support common controls on motifs diversity. We hypothesize that strong interspecific competition and optimized signaling among congeners sustain maximized subdivision in the pigmentation pattern space.

## Pigmentation patterns are evolutionarily labile, yet spatially conserved

Species richness and phylogenetic diversity can vary dramatically between biogeographic regions, suggesting that clade-specific biogeographic histories may have shaped variation in pigmentation diversity between ecoregions. However, we found no significant variation in pigmentation pattern diversity among the biogeographic regions for any fish family (np-MANOVA: surgeonfishes, *p*=0.94; butterflyfishes, *p*=0.47; snappers, *p*=0.18; goatfishes, *p*=0.95; angelfishes, *p*=0.08; damselfishes, *p*=0.06; **Extended Data Table 4**). These results indicate that species groups from each ecoregion occupy the same hypervolume in the pigmentation pattern space (**Extended Data Figure 1**), challenging the hypothesis that pigmentation pattern disparity is partitioned between biogeographic regions. Instead, our results reveal that fish pigmentation pattern diversity is remarkably conserved among biogeographic regions on a global scale.

The consistency of pigmentation pattern diversity can be a consequence of lineages with distinct pigmentation patterns colonizing different biogeographic regions over time, or the repeated divergence of pattern motifs between close evolutionary relatives in sympatry, resulting in convergence between distantly related lineages. To differentiate between these scenarios, we leveraged time-calibrated molecular phylogenies to test the statistical dependence between pigmentation patterns and the phylogeny of each fish family (i.e. phylogenetic signal) using the multidimensional equivalent of Blomberg’s K^28^. In all cases, values of K were close to 0 (Acanthuridae: Kmult=0.11, *p*<0.01; Chaetodontidae: Kmult=0.10, *p*<0.01; Lutjanidae: Kmult=0.22; *p*<0.01; Mullidae: Kmult=0.08, *p*<0.01; Pomacanthidae: Kmult=0.05, *p*<0.01; Pomacentridae: Kmult=0.06, *p*<0.01), indicating a very low degree of phylogenetic relatedness in pigmentation patterns. These low K values demonstrate that closely related species exhibit more divergent pigmentation patterns than would be expected under a null Brownian motion model, with significant *p*-values in all cases revealing that diversification of pigmentation patterns is non-random. These results suggest the lack of deviation in pigmentation pattern diversity among biogeographic regions is likely driven by the repeated divergence of pattern motifs between closely-related lineages.

Our findings resolve the seemingly paradoxical relationship between high trait lability and regional conservatism. Coral reef fishes possess the largest array of pigment cell types among vertebrates^5^. It is possible that regional conservatism may reflect underlying molecular and cellular constraints on the generation of new motifs (e.g., pattern formation mechanisms, pigment cell diversity, physical pigment properties, etc^7,29,30^). For example, labyrinthine motifs have been proposed to originate from interspecific hybridization between spotted species^6^. Likewise, new pigmentation patterns can be produced by a simple modification of developmental mechanisms^31,32^. Our findings, alongside studies on the molecular basis of pigmentation^6,7,33^, suggest that a limited set of developmental mechanisms and associated genes drive the evolution of pigmentation patterns, leading to recurrent phenotypic convergences under various selective pressures. This constrained evolutionary process may reflect a broader principle of phenotypic evolution, where natural selection operates on a limited set of developmental processes, resulting in recurrent patterns of convergence across diverse taxa.

## Mode and tempo of pigmentation pattern evolution

Variation in pigmentation patterns can influence the divergence and maintenance of reef-fish species boundaries^20,34^, suggesting that motifs diversity may disproportionately diverge among closely related taxa and lead to accelerated diversification rates among tip-ward lineages. We tested this hypothesis through a comparison of five evolutionary models: 1) Brownian motion (BM) of stochastic diffusion (modeling gradual evolution); 2) the ψ (psi) model of mixed gradual and speciational evolution^35^; 3) the Ornstein-Uhlenbeck (OU) model that combines stochastic diffusion with a pull towards a central “optimum” value^36^; 4) an early burst (EB) model that describes an exponential decrease in evolution as is often expected in adaptive radiation, and the accelerating (AC) model where evolutionary rates increase exponentially over time. Using sample-size corrected Akaike’s Information Criterion (AICc) for model selection, we found strong support for the AC and OU models as the best fitting models for the majority of principal component (PC) axes, capturing 90% and 100% of the fitted PCs, respectively (**Figure 2, Extended Data Table 5**). The strong support for the AC model indicates that pigmentation pattern diversification has accelerated during the recent evolution of reef fish families.

**Figure 2.**
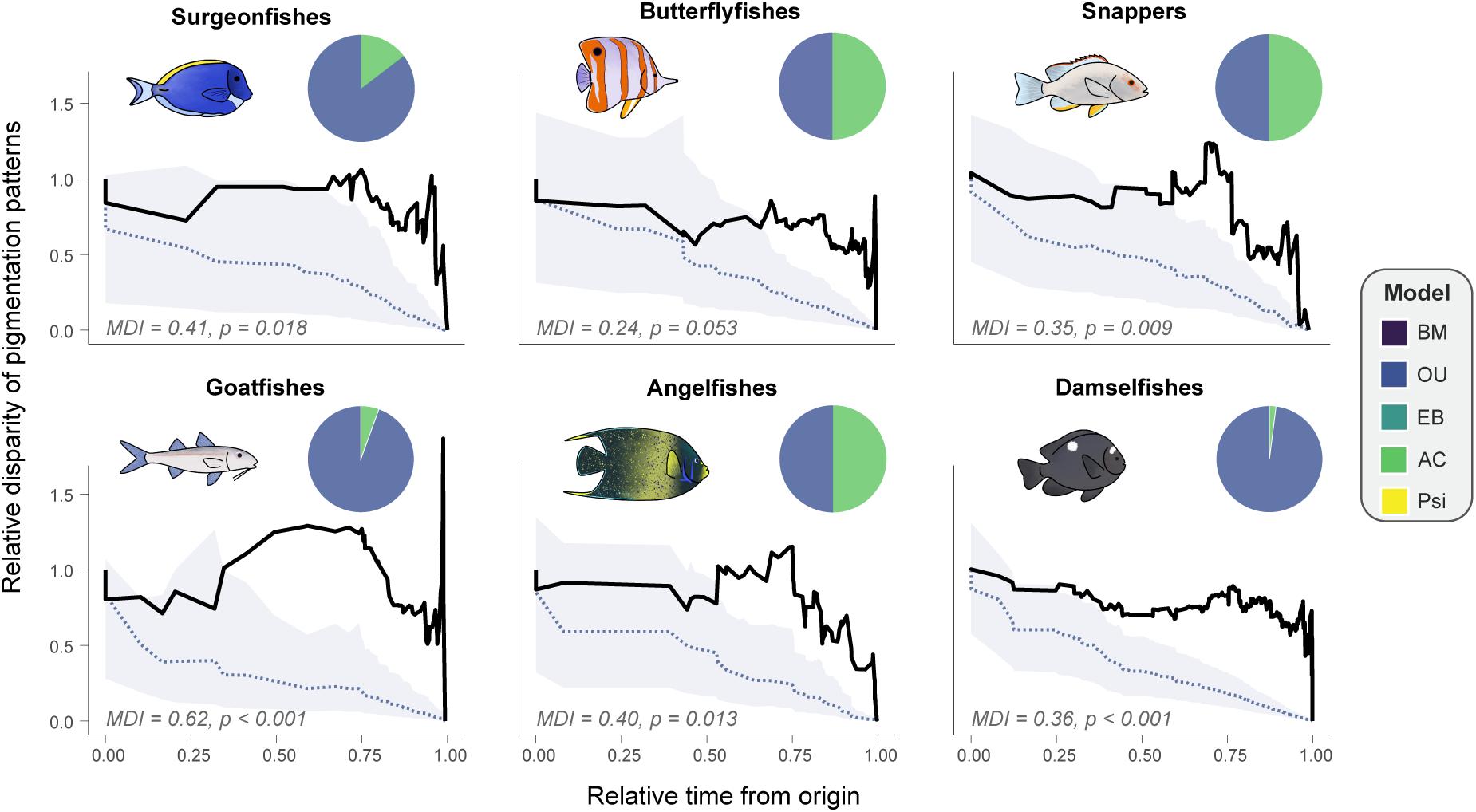
*Disparity through time (DTT) plots* illustrating the evolution of subclade disparity across the phylogeny of each fish family. Higher relative disparity values (y-axis) correspond to a greater average volume of pigmentation pattern space occupied by subclades relative to the disparity of the entire family. The solid line represents the observed disparity calculated for each clade, while dashed line is median expected disparity under a null Brownian motion model based on simulations. Across all fish families, subclade disparity was significantly higher than expected under BM, with sharp increases of within subclade disparity toward the present, that suggest the potential for high motif convergence between subclades. Inset *Pie charts* display the distribution of the Akaike information criterion weights (AICw) from trait evolution model (BM, OU, EB, AC and psi) comparisons. Generally, BM, EB and psi models support values were too low to be discernable on pie charts. Above analyses reflect model comparisons for PC1. The analyses of PCs 2-3 are provided in Extended Data Figure 2 and Extended Data Table 5.

It is possible that varying environmental conditions and competitive interactions could have promoted either early pulses or gradual accumulations of motif diversity within fish assemblages over time. However, our results suggest this to not be the case. The poor fit of the EB and BM models allowed us to reject hypotheses of decelerating evolutionary rates or constant diversification of pigmentation patterns. In contrast, we found strong support for the fit of the single-optimum OU model (OU1) to many PC scores (**Figure 2, Extended Data Table 5 & Extended Data Figure 3**). The strong support for the OU1 model indicates that while pigmentation patterns evolve rapidly, they do so around a constrained set of motifs. This result is concordant with our analyses of the occupation of pigmentation pattern spaces (**Extended Data Figure 1**) and aligns with the high level of homoplasy observed in animal coloration, where distantly related species often share similar pigmentation patterns. These results support expectations of iterative evolution, where pigmentation motifs repeatedly emerge from a central set of patterns. Our finding of this rapid, yet bounded, diversification pattern suggests the possibility that this diversification may be a feature of pigmentation motif evolution, as the pattern of homoplasy we observe in coral reef fishes closely mirrors those in other vertebrates, such as the plumages of birds^37,38^.

The bounded trait space supported by our analyses raises the possibility that pigmentation motifs may lose evolvability due to developmental constraints possibility limiting reversals or transitions between specific motifs. However, the accelerated evolutionary rates detected in our study suggest that pigmentation patterns remain highly labile. We estimated a very short phylogenetic half-life [i.e. the time to evolve half the distance between the root state and the selective optimum value (t1/2 = ln(2)/α)] for motifs, with a median value of 1.5 million years across fish families (**Extended Data Table 5**). This half-life is extremely rapid when contrasted with morphological traits in fishes (median half-life of 37.5 million years across 13 morphological traits^39^). Furthermore, the best-fit model for PC2 in butterflyfishes was the λφϑ model (**Extended Data Figure 2**), illustrating that speciation and aspects of pigmentation pattern evolution are also closely linked. Collectively, these results underscore that pigmentation pattern diversification reflects highly rapid evolution within a bounded space, that is likely linked to rapidly shifting ecological or other selective pressures.

The observed pattern of rapid, yet bounded, evolution in pigmentation motifs may simply reflect a stochastic distribution of motif diversity across the phylogeny with no general phylogenetic structuring of trait disparity. We tested this possibility by assessing the partitioning of trait disparity among subclades over time through the quantification of morphological disparity index (MDI) values in an analysis of subclade disparity through time (DTT). In all cases, DTT plots of PC axes 1 to 3 demonstrated a sharp increase within subclade pigmentation pattern disparity in recent evolutionary history, with positive MDI values indicating a significant departure above median Brownian motion expectations (**Figure 2, Extended Data Figure 3**). This pattern of recent subclade disparity being partitioned within versus between subclades, provides strong support for the hypothesis that pigmentation pattern evolution has been characterized by consistent and repeated divergences between closely related taxa that culminate in convergences in pigmentation patterns among distantly related lineages. Additionally, stochastic mapping of motif shifts over time (**Figure 3** **& Extended Data Figure 3)** further support the results of the DTT analysis, revealing most changes in motif patterns occurring geologically recently (within the last 5-10 MYA). These discrete trait analyses (**Figure 3**) closely align with results from continuous trait models (**Figure 2** **& Extended Data Table 5**), and support that the conclusion that pigmentation pattern diversity of extant coral reef fishes evolved recently and rapidly.

**Figure 3.**
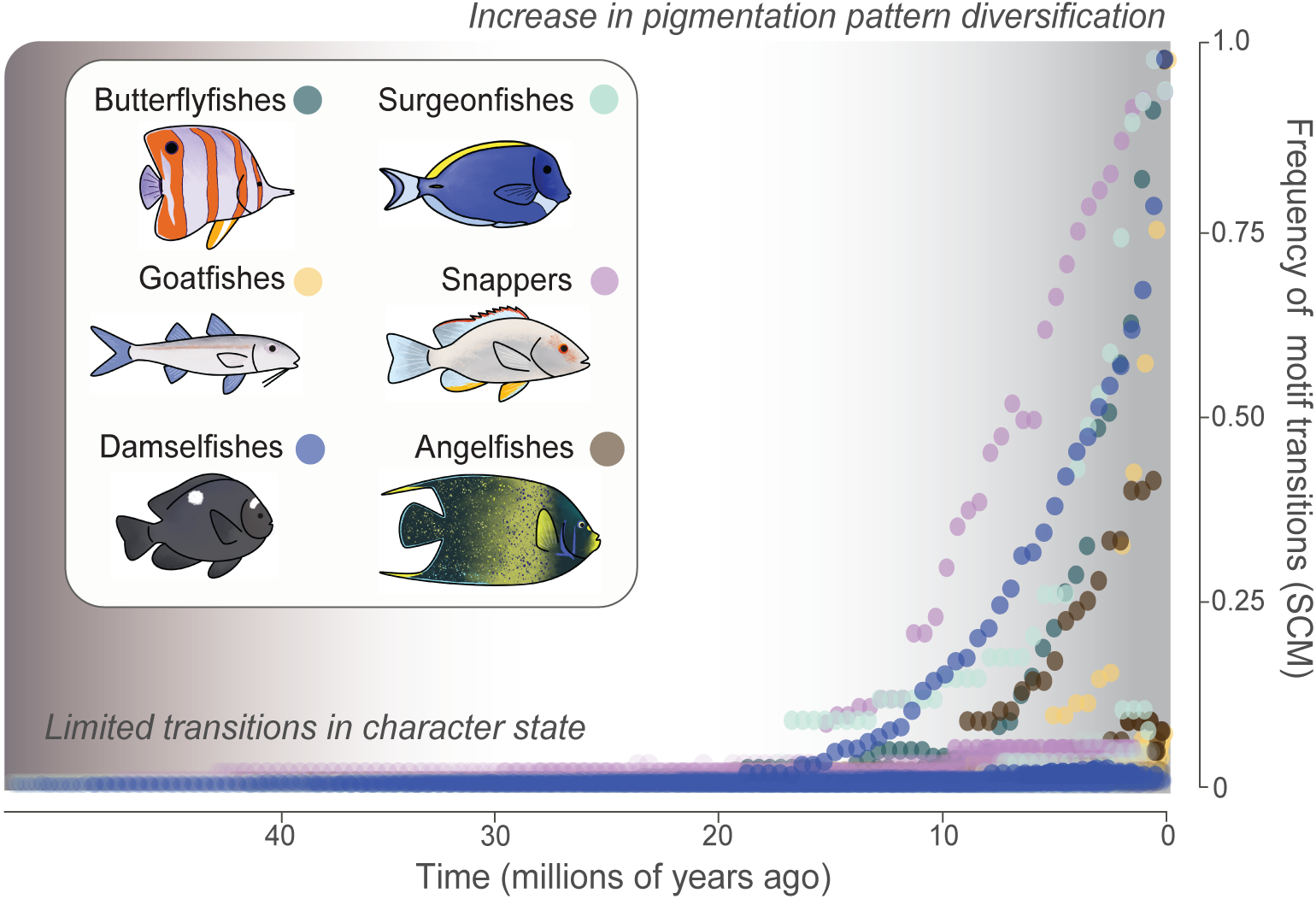
Shifts in motifs over time across all studied fish families. The y-axis provides the frequency of all transitions between presence and absence of each motif across the stochastic character maps (SCM). Circles are the number of transitions across the SCM for every time point (node), which are standardized by the number of lineages present at that time. Circles are colored to correspond to each reef fish family.

Adaptive functions like visual communication, camouflage, mimicry, and aposematism, shaped by both sexual and natural selection^40,41^, likely impose trade-offs that can bound the limits of phenotypic diversity between clades of organisms. However, the strikingly parallel diversification dynamics we observed across all studied reef fish families – which vary dramatically in their ecology and morphology – suggest a strong developmental influence on pigmentation pattern diversity. Cellular mechanisms involved in pigmentation pattern formation coupled with the modularity of pigmentation pattern expression^5^ likely generate motif patchworks across the body. These motifs, in turn, likely result in the recurrent evolution of pigmentation patterns between divergent taxa. This hypothesis parallels pathways that have been suggested for bird plumage^37,42^ and possibly also promoted convergence in reef fish. The rapid and repetitive evolution of pigmentation patterns in reef fishes suggests these visually striking traits are likely ephemeral on a macroevolutionary scale, appearing and disappearing several times throughout evolutionary history. Further research is certainly needed to clarify the developmental and genetic pathways that facilitate this iterative evolutionary process that has given rise to some of the most colorful assemblages of vertebrates on our planet.

## Conclusion

Our findings illuminate factors that explain color pattern diversity in living coral reef fishes, revealing that recent speciation events have driven high levels of pigmentation pattern disparity within subclades. This recent and rapid diversification contrasts with the diversity dynamics of ecomorphological traits like head, fins, and body morphology, in which trait disparity is often partitioned between major subclades^43–46^. However, this does not imply that pigmentation pattern diversity did not exist in earlier geologic periods. On the contrary, motifs similar to those found today, such as spots and stripes, appear on fossilized fish from the Eocene^34^, suggesting color pattern motifs to be both evolutionarily labile and ephemeral. These observations parallel findings in birds, where rapid plumage evolution also displays high evolutionary lability and homoplasy^37,42^.

Mechanistically, studies in model teleosts such as zebrafish have found pigment cell signaling and communication dynamics as key to generating diverse pigmentation patterns^7,31,47^. When considering the wide array of pigment cell types and color polymorphisms in coral reef fishes^5,48,49^, these findings highlight the rich molecular and cellular substrate of pigmentation diversity upon which selection acts. Additionally, processes such as introgression^50,51^ and hybridization, especially in visually striking groups like butterflyfishes, angelfishes, and clownfishes^52,53^, possibly further fuel rapid and repetitive motif evolution. Future research investigating the role of hybridization and developmental pathways in pigmentation pattern formation will be crucial for understanding the evolutionary dynamics of these traits. Moreover, considering how robust these labile evolutionary traits are to changes in species ranges, water turbidity, and habitat in response to environmental shifts over the next century remains an urgent priority.

## Methods

### Data collection

Fish images from illustrated scientific books and from public databases (e.g., FishBase: https://www.fishbase.org) were used to categorize the presence or absence of pattern motifs. Our dataset includes a total of 918 species: 83 acanthurids, 133 chaetodontids, 128 lutjanids, 80 mullids, 92 pomacanthids and 402 pomacentrids, representing a minimum of 75% of the extant diversity of each fish family (**Extended Data Table 1 & Extended Data Methods**). We aimed to use a minimum of three images per species. Pigmentation patterns were described at the adult stage only. If sexual dimorphism is present, we only quantified the pigmentation pattern observed in males.

Five ecoregions mainly aligned with major ocean basins were delineated, based on the methodology of Kulbicki et al.^13^. These ecoregions are the Atlantic Ocean (including the Mediterranean Sea), the Western Indian Ocean (including the Red Sea), the Central Indo-Pacific (extending from the edge of India to the northern edge of New-Zealand), the Central Pacific and the Tropical East Pacific. Geographical distribution patterns of each species were extracted from FishBase^24^, which provides a validation of previously published assigned areas (i.e. miss-identification or changes in species taxonomy), and GBIF^25^. A species was considered to occupy an ecoregion when at least five observation points were displayed within it. Species having a widespread geographical distribution encompassing different ecoregions were coded as occupying each of these related ecoregions.

### Annotation of fish pigmentation patterns

Fish images were subjected to pattern annotation by the same three observers: L. Mittelheiser, J.R. Hodge and L. Moulin. This task was triple checked independently by each observer. To describe the pigmentation pattern of each fish species, we targeted motifs commonly used in publications related to this topic^6^. Pigmentation patterns were initially annotated independently on the three main areas of the body (head, trunk and tail) and then merged into a single “total” dataset. We formulated six classes of patterns: the number of colors (one or more), color separation (three possible types: H-sep - different background colors are separated following the body axis ; V-sep - different background colors are separated perpendicularly to the body axis ; O-sep - different background colors are separated with an orientation deviating from the body axis from around 20° to 70°), stripe patterns (five possible types: H-stripes - horizontal stripe following the body axis ; V-stripe - vertical stripe oriented perpendicularly to the body axis ; O-stripes - oblique stripe deviating from the body axis from about 20° to 70° ; Marbling: irregular, elongated and connected patterns without a directional trend ; and Honeycomb: juxtaposed hexagonal patterns), blotch patterns (two types : Regular blotches - large and irregular markings ; Saddle - blotch located on dorsal part ahead of the caudal peduncle), type of colored structures (two structures: Mouth - mouth has a different color than head’s background color ; Scalpel - scalpel has a different color from body’s background color), and other patterns (four possible types: Dots - small and rounded area of color ; Reticulations - net-like pattern ; Eyespot - large dot surrounded by a ring of a different color ; Edging - structure is outlined with a color different from the structure’s background color). Each of these traits was coded as present (1) or absent (0) in studied taxa. Illustrations and detailed descriptions of every motif are provided in the **Extended Data Methods**. The diversity of pigmentation patterns varied across the studied families. Following our objectives to best describe the diversity of phenotype within each taxon, the fish families were studied separately in the downstream analyses.

### Quantitative analysis of pigmentation patterns

All analyses were conducted in R version 4.3.0^54^. To summarize the diversity of motifs and to create a pigmentation pattern space, we calculated the distance between all pairs of species using Gower’s metric. Then, we applied a Principal Coordinates Analyses (PCoA, also called principal axes for simplicity) on this matrix^55^. Following the framework of Maire and colleagues^56^, we identified the lowest number of dimensions to capture phenotypic diversity without losing information by performing a quality test using the function *quality.fspaces()* in the R-package mFD^57^ (version 1.0.7). Accordingly, we used species’ coordinates on the first eight principal axes of the PCoA for Acanthuridae and Chaetodontidae; on the first seven principal axes for Pomacanthidae and Pomacentridae; and on the first six principal axes for Lutjanidae and Mullidae. Scatter plots illustrating the dispersion of species in the phenotypic space were produced using the function *funct.space.plot()* from the R-package mFD (Extended Data Figure 1).

To explore the occupation of pigmentation pattern space, we used the multifaceted framework of functional ecologists^26,58^ to decompose the phenotypic diversity into three complementary components: *Richness*, *Divergence* and *Evenness*^26^. *Richness* is the total extent of multidimensional space utilized by species. *Divergence* quantifies the distribution of species within the multidimensional space. *Divergence* approaches zero when species are close to the center of gravity of the space (or hypervolume) occupied by the group of species in the phenotypic space; it equals one when species are located on the edges of the space. The higher the value of this index, the more species are differentiated in the multidimensional space. *Evenness* characterizes regularity in the distribution of species along the shortest tree linking all of them. *Evenness* approaches zero when species are packed within a small region of the multidimensional space, and it equals one for an even distribution of species in the space. High values of this index indicate a limitation in phenotypic similarity. The three indices were computed using the *function alpha.fd.multidim()* from the R-package mFD.

Variation in the diversity of pigmentation patterns among fish assemblages from the five ecoregions was assessed by using a non-parametric MANOVA (np-MANOVA)^59^, performed with the function *procD.lm()* using a randomized residual permutation procedure (RRPP) and 10,000 permutations from the R-package *geomorph*^60^ (version 4.0.8). As for the calculation of the three functional indices, we applied np-MANOVA analyses by using the number of principal axes suggested by quality tests.

One species can be present in multiple ecoregions. Thus, we first calculated *Richness*, *Divergence* and *Evenness* indices and performed np-MANOVA by assigning widespread species in all the regions where encountered. However, the presence of species in multiple ecoregions could have a homogenizing effect on any differences among regions. Accordingly, *Richness*, *Divergence* and *Evenness* indices as well as np-MANOVAs were also computed using re-sampling techniques in which any species present in more than one ecoregion was randomly assigned to a single ecoregion. Re-sampling was performed with 1,000 iterations and the median values and standard deviations of functional indices and np-MANOVA were retained (see **Extended Data Methods** for further details).

### Phylogenetic comparative analyses

Beyond exploring pigmentation pattern space occupation, we also studied the evolution of pigmentation patterns by using a combination of comparative analyses that account for phylogenetic relatedness. We used published, strongly supported multi-gene, time-calibrated phylogenies. All details concerning the methods of phylogenetic reconstruction and time calibration are provided in these works: see Sorenson et al.^61^ for Acanthuridae, Hodge et al.^62^ for Chaetodontidae, Rincon-Sandoval et al.^63^ for Lutjanidae, Nash et al.^46^ for Mullidae, Baraf et al.^64^ for Pomacanthidae and McCord et al.^65^ for Pomacentridae.

Phylogenetic signal is a measure of the statistical dependence among species’ trait values on their phylogenetic relationships. It can be used to describe the conservatism or the evolutionary lability of traits across phylogenetic histories^66^. Here, we applied the multivariate version of the Blomberg’s K (Kmult) available in the R-package *geomorph* (function *physignal*) to the number of principal axes of the pigmentation pattern space retained by the quality tests. Similar to the interpretation of the univariate Blomberg’s K^28,67^, Kmult = 1 indicates strong phylogenetic signal that perfectly follows Brownian motion. A Kmult value > 1 means that closely related species trait values are more similar than expected under a Brownian motion model, whereas a value < 1 suggests a greater lability of trait values and a departure from a strong phylogenetic signal.

We tested the hypothesis that pigmentation patterns diversified recently in reef fish families by comparing the fit of five models of trait evolution. We fitted a single rate (σ^2^) Brownian motion (BM) model, an Ornstein-Uhlenbeck model with a single optimum (θ) across the entire tree (OU1), a ψ (psi) model of mixed gradual and speciational evolution, an early burst (EB) model where the rate of evolution decreases exponentially through time, and an accelerating (AC) model where the rate of evolution increases exponentially through time. Modeling of continuous trait evolution was conducted using the function *transformPhylo.ML()* in the R-package *motmot* ^68,69^ (version 2.1.3). For these comparative analyses, we fitted models separately on the first three principal axes to optimize convergence on reliable solutions. Models were compared using AICc (Akaike’s information criterion, corrected for sample size) and Akaike weights, which balance goodness of fit with model complexity^70^.

We further characterized the tempo of pigmentation pattern diversification using disparity-through-time analysis^71^. This approach computes the average subclade disparity for one trait at each node in the phylogeny and plots these as a function of node age. At the root of the tree, the average subclade disparity is simply the phenotypic disparity of the entire clade and is therefore high. At subsequent nodes, disparity is averaged over the total number of subclades in existence at that time. Under an early burst hypothesis, average subclade disparity is expected to decline rapidly in the early history of the clade as evolutionary rates slow and phenotypic variation becomes partitioned among subclades. Conversely, an average subclade disparity remaining high or even increasing through time signals high phenotypic variation within clades, potentially due to convergence. We also used the morphological disparity index (MDI) of Harmon et al.^71^ to quantify the difference between average subclade disparity through time for our observed dataset and that expected under a null BM model. Negative MDI values indicate lower than expected subclade disparity relative to BM while positive MDI values indicate higher than expected subclade disparity. MDI statistics were computed for the first three principal axes over the first 90% of the time-tree using the R-package *geiger*^72^ (version 2.0.11). We omitted the most recent 10% of the phylogeny in our analysis to avoid spurious MDI estimates due to incomplete sampling of tip species^71^.

Finally, we quantified the dynamics of pigmentation pattern diversification by counting the number of motif shifts over evolutionary time. This was achieved by producing stochastic maps for every single motif using the function *make.simmap()* in the R-package *phytools*^73^ (version 2.1-1). Here, we sampled 10,000 character histories allowing the incorporation of the uncertainty associated with the timing of the transitions between presence and absence of the pattern motif, based on the best fitting model of trait evolution using AIC. For the parametrization of *make.simmap()*, we used the estimated ancestral state and an equal rates transition matrix between states. Data were then summarized by plotting the frequency of motif transitions over time, as well as the distribution of times spent in either character state for each trait across the phylogeny.

## Acknowledgements

We thank L. Moulin for assistance with dataset assembly. We thank C. Petetin for creating the fish images used on every figure.

## Authors contributions

B.F., V.L. and A.D. designed the study. B.F. and A.D. drafted the paper with inputs from L.M., A.G., J.H. and V.L. L.M. collected phenotypic and geographic data. J.H. contributed to phenotyping. B.F., L.M., A.G. and A.D. analysed data. All authors contributed to interpretation and discussion of results.

