## Supplementary materials for "Rapid and repeated evolution of the pigmentation patterns in reef fishes"

**List of contents:**

- Extended Methods:
    1) Summary of taxon sampling.
    2) Description of pigmentation motifs.
    3) Resampling methods for the calculation of functional indices and np-MANOVA.

- Extended Table 1.
- Extended Table 2.
- Extended Table 3.
- Extended Table 4.
- Extended Table 5.

- Extended Figure 1.
- Extended Figure 2.
- Extended Figure 3.

**Extended Methods**

***Summary of taxon sampling and number of used images***

Proportion of extant diversity is based on total number of species given by Eschmeyers Catalog of Fishes (https://www.calacademy.org/scientists/projects/eschmeyers-catalog-of-fishes).

| | Number of studied species | Proportion of extant diversity (%) | Number of images |
|---|---|---|---|
| Acanthuridae | 83 | 98 | 83 |
| Chaetodontidae | 133 | 97 | 288 |
| Lutjanidae | 128 | 96 | 128 |
| Mullidae | 80 | 79 | 80 |
| Pomacanthidae | 92 | 100 | 94 |
| Pomacentridae | 402 | 93 | 1126 |
| **Total** | **918** | | **1799** |

***Description of the 28 traits (motifs) to quantify the patterns of pigmentation***

Hereafter, we provide a short description of each trait used to characterize the pigmentation pattern in fishes. Each trait was reiterated for the head, the trunk, and the tail. We defined the three body areas as followed: the head region extends to the posterior extremity of the opercula; the trunk region begins right after the opercle and stops at the beginning of the caudal peduncle; the tail region initiates at the end of the trunk, encompassing both the caudal peduncle and the caudal fin.

1) *One color*: region is composed of one color in the background.

Head

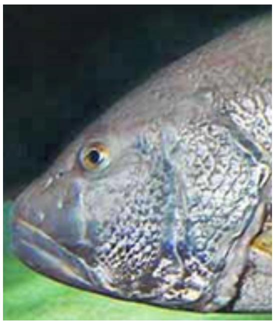

Trunk

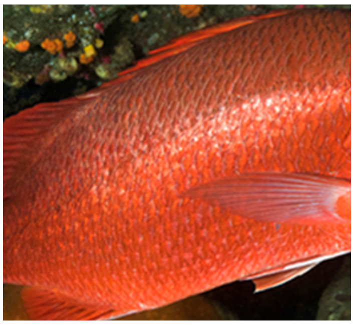

Tail

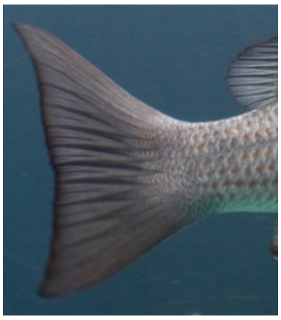

2) *Two or more colors*: region is composed of two or more colors in the background.

Head

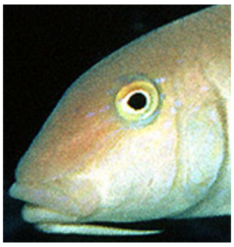

Trunk

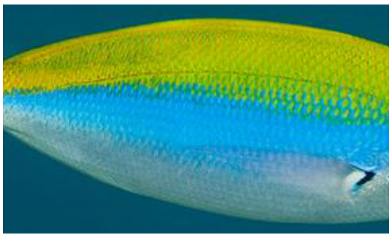

Tail

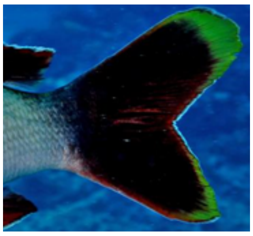

3) *One horizontal stripe*: region exhibits one long, horizontal, straight area of a single color.

Head

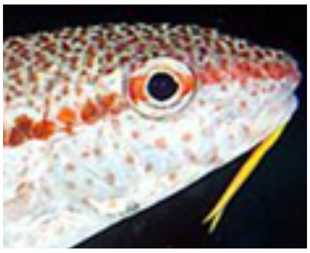

Trunk

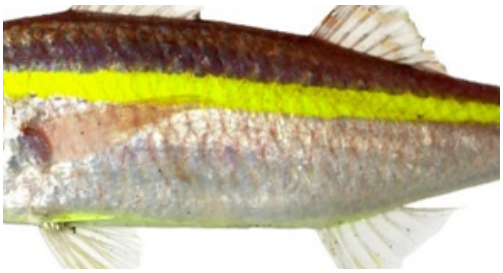

Tail

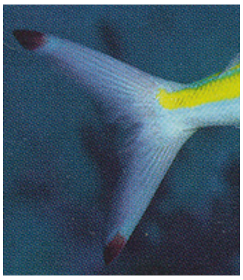

4) *Two or more horizontal stripes*: region exhibits two or more long, horizontal, straight areas of a single color.

Head

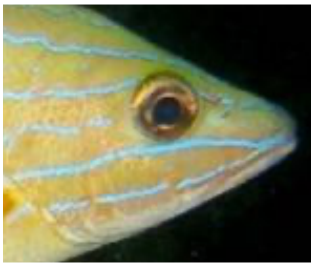

Trunk

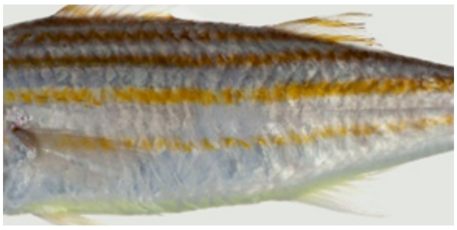

Tail

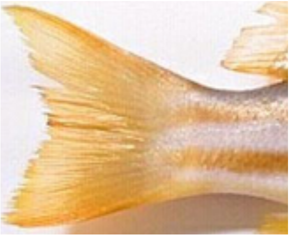

5) *One vertical stripe*: region exhibits one long, vertical, straight area of a single color.

Head

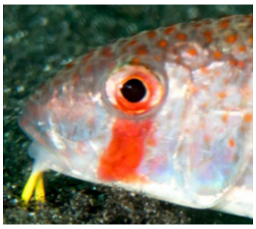

Trunk

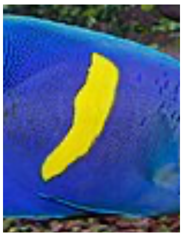

Tail

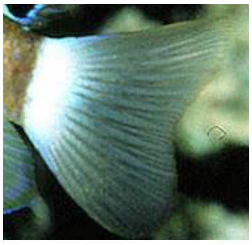

6) *Two or more vertical stripes*: region exhibits two or more long, vertical, straight areas of a single color.

Head

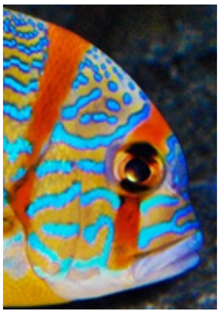

Trunk

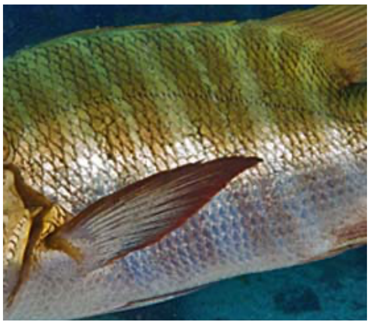

Tail

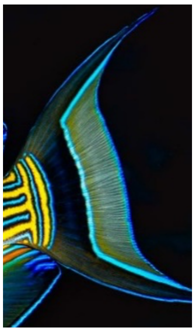

7) *One oblique stripe*: region exhibits one long, oblique, straight area of a single color.

Head

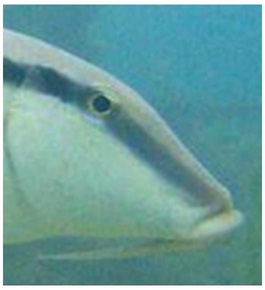

Trunk

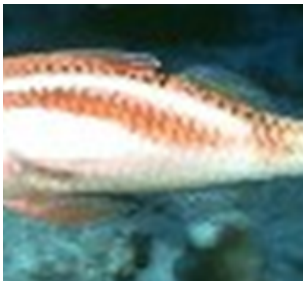

Tail

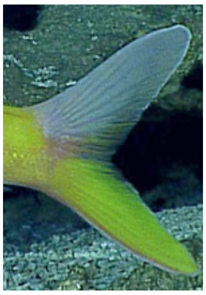

8) *Two or more oblique stripes*: region exhibits two or more long, oblique, straight areas of a single color.

Head

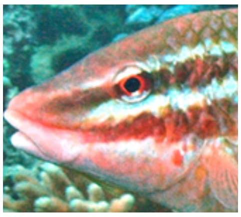

Trunk

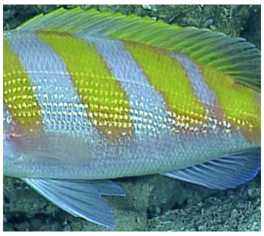

Tail

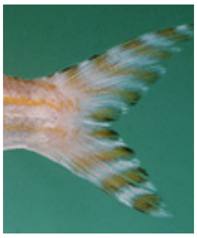

9) *One blotch*: region exhibits a large and irregular patch of a single color.

Head

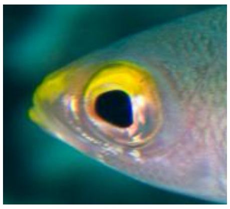

Trunk

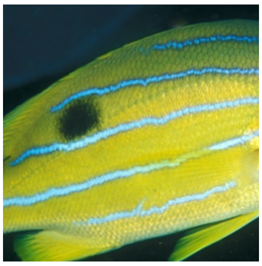

Tail

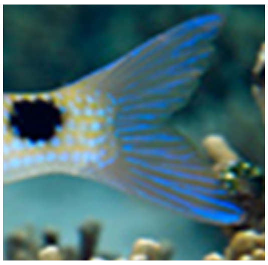

10) *Two or more blotches*: region exhibits two or more large and irregular patches of a single color.

Head

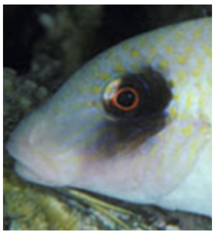

Trunk

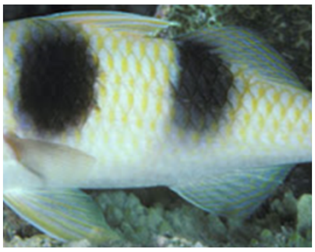

Tail

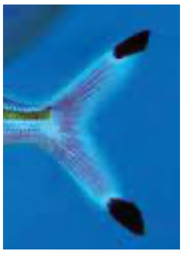

*11) One dot*: region exhibits one small and regularly circular area of a single color.

Head

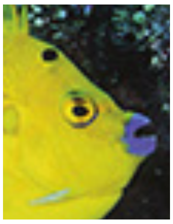

Trunk

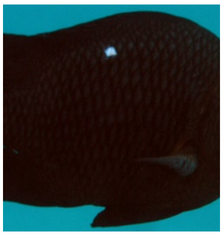

*12) Two or more dots*: region exhibits two or more small and regularly circular areas of a single color.

Head

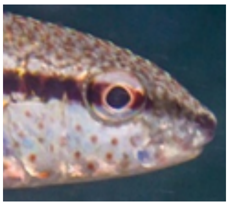

Trunk

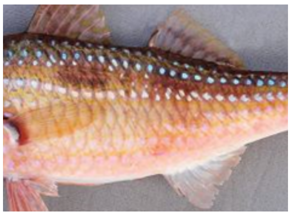

Tail

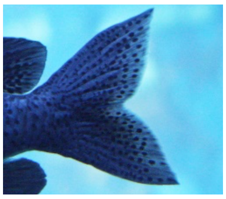

*13) Eyespot*: region exhibits a large regularly area of a single color surrounded by a ring of a different color.

Head

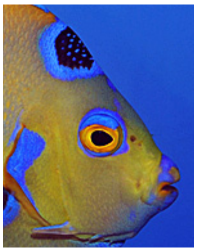

Trunk

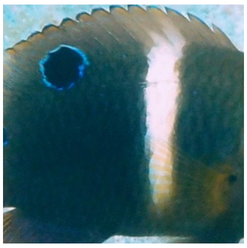

Tail

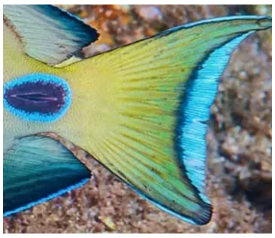

*14) Marbling*: region exhibits irregular, elongated, connected areas of a single color.

Head

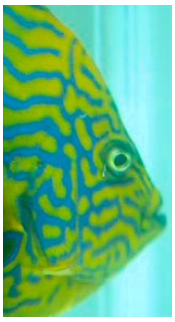

Trunk

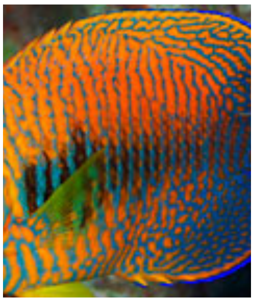

Tail

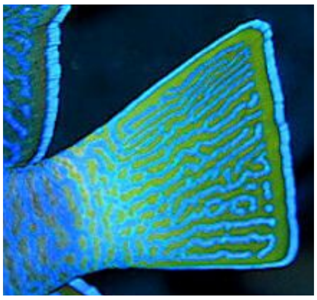

15) *Eye stripe*: Eye exhibits a long, straight area of a single color.

Horizontal

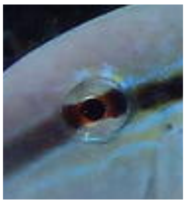

Vertical

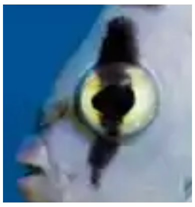

16) *Color separation orientation on the trunk*: for species exhibiting two or more colors in the background of their trunk, the direction in which these colors are separated. When there are more than two colors, there can be multiple orientation for the same region.

Horizontal

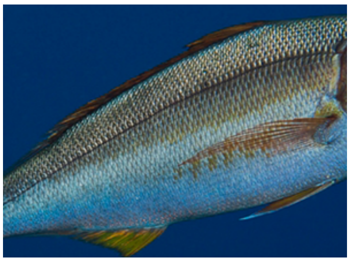

Vertical

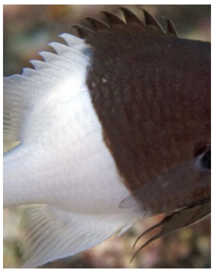

Oblique (Yellow and blue)

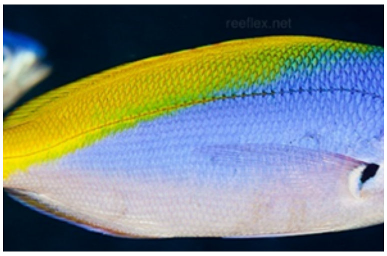


17) *Saddle*: large and irregular area of a single color extending from the sides over the dorsal surface, ahead of the caudal peduncle.

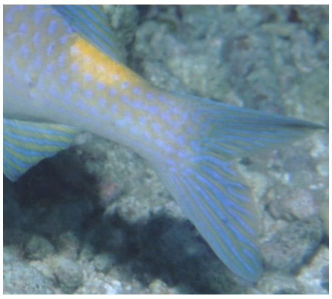

18) *Stripes on dorsal fin*: dorsal fin exhibits one or more long, straight areas of a single color, whatever the orientation.

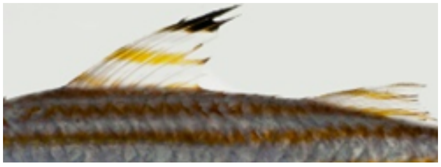

19) *Blotches on dorsal fin*: dorsal fin exhibits one or more large and irregular areas of a single color.

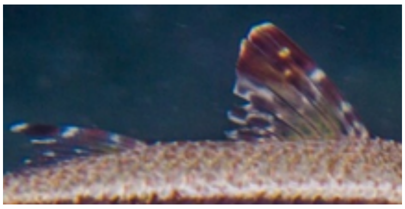

20) *Colored mouth*: the mouth exhibits a different color than the head's background color.

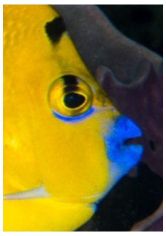

21) *Encircled eye*: the eye is surrounded by a circle with a different color than the head's background color.

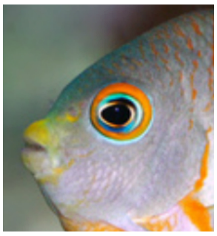

22) *Operculum border*: a thin area of a single color, different from the head's background color, following the border of the opercular bone and/or the preopercular bone.

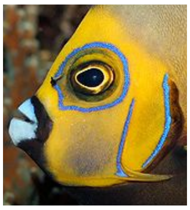

23) *Blotch at the pectoral fin insertion*: an irregular area of a single color located at the pectoral fin insertion.

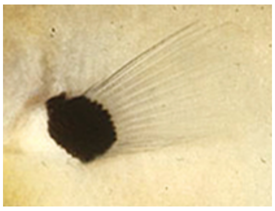

24) *Caudal fin border*: a thin area of a single color, different from the caudal fin background color, follow the border of the caudal fin.

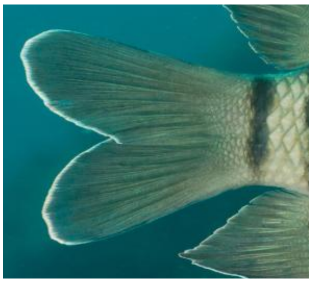

25) *Blotches in the eye*: an irregular area of a single color, different from the background color of the eye.

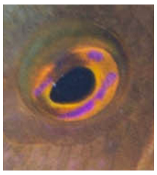

26) *Honeycomb pattern*: area exhibits hexagonal patterns, looking like honeycomb.

Head

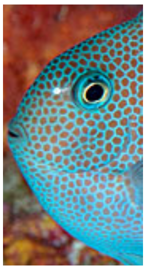

Trunk

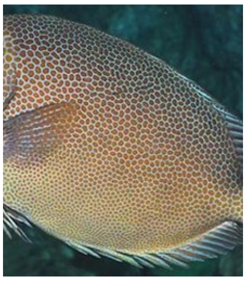

Tail

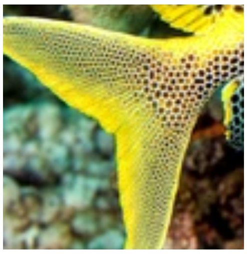

27) *Colored scalpel* (trait for surgeonfish only): scalpel(s) color is different from the background body color surrounding it.

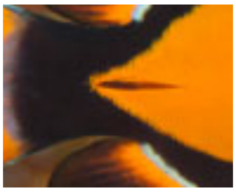

28) *Blotch around the scalpel(s)* (trait for surgeonfish only): an irregular area of a single color, different from the background color of the body, surrounding scalpels

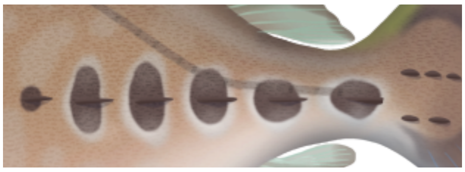

***Resampling methods for functional indices and np-MANOVA***

Out of the 918 species included in our dataset, 516 species were present in more than one ecoregion (Acanthuridae: 54 species in more than one ecoregion; Chaetodontidae: 46 species; Lutjanidae: 62 species; Mullidae: 38 species; Pomacanthidae 18 species; Pomacentridae 99 species). To avoid a potential homogenization of pigmentation patterns across ecoregions in quantitative analyses due to the presence of a given species in several ecoregions, re-sampling methods were used to compute functional indices (motifs *Richness*, *Divergence*, and *Evenness*) and np-MANOVA comparing variations in pattern diversity across ecoregions. All species in the dataset were forced to be assigned to a single ecoregion so that any species present in several ecoregions was randomly assigned to one of the ecoregions where it is present and labeled as absent in other ecoregions.

Re-sampling was performed 1,000 times for each family. Functional indices and np-MANOVA were computed for each iteration. The median value and standard deviations of each index and of the F-values and *P*-values of np-MANOVA tests were retained and are presented in the results.

The *alpha.fd.multidim* function (*mFD* R-package) used to compute functional indices requires the number of observations (here, species) to be larger than the number of variables (here, PC axes) used in the analysis. In iterations where re-sampling led to a number of species smaller than the number of PC axes retained for a given family, duplication of a few species in two ecoregions was allowed until the minimum number of species required by the function was reached. Note that this was not a requirement for the np-MANOVA, so species were not duplicated for this test.

**Extended Table 1.** Provided as a .xlsx file (Extended_Data_Table1.xlsx). List of the studied species (918) in surgeonfishes (Acanthuridae), butterflyfishes (Chaetodontidae), snappers (Lutjanidae), goatfishes (Mullidae), angelfishes (Pomacanthidae), and damselfishes (Pomacentridae). For clarity, families are distributed on different sheets. The coding of ecoregions (biogeographic data) and motifs (raw data for the quantification of pigmentation patterns) are provided.

**Extended Table 2.** Motif *Richness* (Rich), *Divergence* (Div), and *Evenness* (Eve) in each ecoregion (CIP: Central Indo-Pacific; WI: Western Indian; CP: Central Pacific; A: Atlantic; TEP: Tropical Eastern Pacific) for the six fish families. Indices were calculated by (1) assigning widespread species in all the regions where encountered (values in italic) and by (2) resampling methods (1,000 iterations) for which median values (with standard deviations) are provided.

| Family | Index | CIP | WI | CP | A | TEP |
|---|---|---|---|---|---|---|
| Acanthuridae | Rich | *0.61*<br>0.08 (0.04) | *0.45*<br>0.05 (0.03) | *0.49*<br>0.02 (0.02) | *$3x10^{-5}$*<br>$3x10^{-5}$ | *$3x10^{-5}$*<br>$3x10^{-5}$ |
| (surgeonfishes) | Div | *0.77*<br>0.76 (0.01) | *0.79*<br>0.80 (0.02) | *0.78*<br>0.78 (0.02) | *0.87*<br>0.87 | *0.83*<br>0.95 |
| | Eve | *0.85*<br>0.86 (0.02) | *0.87*<br>0.87 (0.02) | *0.85*<br>0.85 (0.02) | *0.95*<br>0.83 | *0.88*<br>0.88 |
| Chaetodontidae | Rich | *0.68*<br>0.40 (0.06) | *0.35*<br>0.25 (0.05) | *0.47*<br>0.13 (0.04) | *0.02*<br>0.02 | *$6x10^{-4}$*<br>$6x10^{-4}$ |
| (butterflyfishes) | Div | *0.81*<br>0.81 (0.01) | *0.83*<br>0.83 (0.01) | *0.82*<br>0.84 (0.01) | *0.83*<br>0.83 | *0.74*<br>0.74 |
| | Eve | *0.81*<br>0.80 (0.01) | *0.82*<br>0.86 (0.01) | *0.86*<br>0.85 (0.02) | *0.83*<br>0.83 | *0.82*<br>0.82 |
| Lutjanidae | Rich | *0.82*<br>0.24 (0.08) | *0.46*<br>0.05 (0.03) | *0.19*<br>$4x10^{-3}$ (0.01) | *0.11*<br>0.01 | *0.01*<br>$2x10^{-4}$ ($1x10^{-5}$) |
| (snappers) | Div | *0.72*<br>0.70 (0.01) | *0.71*<br>0.72 (0.02) | *0.72*<br>0.74 (0.04) | *0.73*<br>0.72 | *0.78*<br>0.83 |
| | Eve | *0.71*<br>0.75 (0.02) | *0.75*<br>0.79 (0.03) | *0.74*<br>0.79 (0.05) | *0.67*<br>0.71 | *0.83*<br>0.84 |
| Mullidae | Rich | *0.52*<br>0.14 (0.05) | *0.31*<br>0.09 (0.04) | *0.37*<br>0.05 (0.04) | *$3x10^{-4}$*<br>$2x10^{-4}$ ($9x10^{-5}$) | *$8x10^{-5}$*<br>$8x10^{-5}$ |
| (goatfishes) | Div | *0.79*<br>0.79 (0.02) | *0.79*<br>0.79 (0.01) | *0.76*<br>0.74 (0.02) | *0.78*<br>0.79 (0.02) | *0.74*<br>0.74 |
| | Eve | *0.83*<br>0.81 (0.02) | *0.84*<br>0.84 (0.02) | *0.83*<br>0.82 (0.03) | *0.81*<br>0.84 (0.03) | *0.85*<br>0.85 |
| Pomacanthidae | Rich | *0.77*<br>0.65 (0.05) | *0.36*<br>0.23 (0.06) | *0.44*<br>0.25 (0.08) | *0.09*<br>0.08 (0.01) | *$8x10^{-5}$*<br>$8x10^{-5}$ |
| (angelfishes) | Div | *0.80*<br>0.78 (0.02) | *0.80*<br>0.78 (0.01) | *0.78*<br>0.75 (0.03) | *0.73*<br>0.73 | *0.65*<br>0.65 |
| | Eve | *0.79*<br>0.79 (0.01) | *0.78*<br>0.79 (0.02) | *0.77*<br>0.78 (0.03) | *0.69*<br>0.69 (0.02) | *0.67*<br>0.67 |
| Pomacentridae | Rich | *0.75*<br>0.52 (0.05) | *0.19*<br>0.10 (0.02) | *0.15*<br>0.03 (0.01) | *$3x10^{-3}$*<br>$3x10^{-3}$ | *$9x10^{-4}$*<br>$9x10^{-4}$ |
| (damselfishes) | Div | *0.77*<br>0.76 ($4x10^{-3}$) | *0.74*<br>0.73 (0.01) | *0.72*<br>0.72 (0.01) | *0.76*<br>0.76 | *0.74*<br>0.74 |
| | Eve | *0.68*<br>0.69 (0.01) | *0.70*<br>0.71 (0.02) | *0.68*<br>0.70 (0.02) | *0.63*<br>0.63 | *0.70*<br>0.70 |

Note: the values of motifs richness calculated by the two methods can vary for the three most-species rich regions: CIP, WI and CP. These differences are driven by a decrease of the number of species included in each region once working with resampling. Conversely, this is not observed for regions A and TEP because we must keep at least as many species as PCs to conduct calculations. We were therefore obliged to keep all the species of these regions at each resampling. Here is an example with Acanthuridae: between the full model (without resampling) and with resampling, we go from 66 species to 34 in CIP, while for TEP we are at 8 species in full and we stay at 8 species for resampling (constraint where number of species in each ecoregion > number of PCs). Accordingly, the richness decreases strongly for CIP but not for TEP.

**Extended Table 3.** Results from correlations between species richness and functional indices (motif *Richness*, *Divergence*, and *Evenness*) across ecoregions for each family based on median values from re-sampling. *R*: Pearson's R coefficient of correlation, *P*: p-value. Significant *P*-values are indicated in bold.

| **Family** | **Index** | **R** | ***P*** |
|---|---|---|---|
| Acanthuridae (surgeonfishes) | Richness | 0.98 | **0.0039** |
| | Divergence | -0.9 | **0.0390** |
| | Evenness | -0.66 | 0.2300 |
| Chaetodontidae (butterflyfishes) | Richness | 0.97 | **0.0060** |
| | Divergence | 0.38 | 0.5300 |
| | Evenness | -0.44 | 0.4600 |
| Lutjanidae (snappers) | Richness | 0.96 | **0.0097** |
| | Divergence | -0.73 | 0.1600 |
| | Evenness | -0.52 | 0.3700 |
| Mullidae (goatfishes) | Richness | 0.96 | **0.0110** |
| | Divergence | 0.51 | 0.3800 |
| | Evenness | -0.62 | 0.2700 |
| Pomacanthidae (angelfishes) | Richness | 1 | **0.0004** |
| | Divergence | 0.71 | 0.1800 |
| | Evenness | -0.69 | 0.2000 |
| Pomacentridae (damselfishes) | Richness | 0.99 | **0.0020** |
| | Divergence | 0.33 | 0.5900 |
| | Evenness | 0.13 | 0.8400 |

**Extended Table 4.** Results from the np-MANOVA, revealing the absence of phenotypic variation among ecoregions in all studied reef fish families. In the first set of tests (Full), species living in multiple ecoregions were assigned in all regions where encountered. In the second set (Resampling), we used a re-sampling method (1,000 iterations) in which any species present in more than one ecoregion was randomly assigned to a single region. For this set, median (SD) values are provided.

| Testing | Family | df | $R^2$ | F | p-value |
|---|---|---|---|---|---|
| Full | Acanthuridae (surgeonfishes) | 4 | 0.03 | 0.64 | 0.94 |
| | Chaetodontidae (butterflyfishes) | 4 | 0.02 | 1.00 | 0.47 |
| | Lutjanidae (snappers) | 4 | 0.02 | 1.28 | 0.18 |
| | Mullidae (goatfishes) | 4 | 0.02 | 0.58 | 0.95 |
| | Pomacanthidae (angelfishes) | 4 | 0.05 | 1.44 | 0.08 |
| | Pomacentridae (damselfishes) | 4 | 0.01 | 1.53 | 0.06 |
| Resampling | Acanthuridae (surgeonfishes) | 4 | 0.05 (0.01) | 1.04 (0.20) | 0.41 (0.24) |
| | Chaetodontidae (butterflyfishes) | 4 | 0.04 (0.01) | 1.29 (0.21) | 0.14 (0.15) |
| | Lutjanidae (snappers) | 4 | 0.05 (0.01) | 1.56 (0.23) | 0.05 (0.06) |
| | Mullidae (goatfishes) | 4 | 0.04 (0.01) | 0.87 (0.75) | 0.64 (0.24) |
| | Pomacanthidae (angelfishes) | 4 | 0.07 (0.01) | 1.70 (0.18) | 0.02 (0.03) |
| | Pomacentridae (damselfishes) | 4 | 0.02 (0.00) | 1.63 (0.15) | 0.03 (0.03) |

**Extended Table 5.** AICc scores and model specific parameters for models fitted to the scores of the first three axes (PC) of the PCoA summarizing the diversity of pigmentation patterns.

| Family | Trait | Model | AICc | ΔAICc | Parameters |
|---|---|---|---|---|---|
| Acanthuridae (surgeonfishes) | PC1 | **OU** | **-140.12** | **0** | $\alpha$ = 0.47; $\sigma$ = 0.23; root state = -0.002; $t_{1/2}$ = 1.5 My |
| | | **AC** | **-136.59** | **3.53** | $\alpha$ = 0.77; $\sigma_{initial}$ = 1.75E$^{-49}$; root state = -0.0002 |
| | | psi | -122.56 | 17.55 | |
| | | BM | -78.34 | 61.78 | |
| | | EB | -76.13 | 63.99 | |
| | PC2 | **AC** | **-166.33** | **0** | $\alpha$ = 0.41; $\sigma_{initial}$ = 3.57E$^{-13}$; root state = -0.005 |
| | | **OU** | **-166.33** | **0** | $\alpha$ = 0.21; $\sigma$ = 0.002; root state = -0.005; $t_{1/2}$ = 3.3 My |
| | | psi | -143.70 | 22.62 | |
| | | BM | 137.59 | 28.73 | |
| | | EB | -135.39 | 30.94 | |
| | PC3 | **AC** | **-187.56** | **0** | $\alpha$ = 0.28; $\sigma_{initial}$ = 3.20E$^{-10}$; root state = -0.004 |
| | | OU | -182.51 | 5.05 | |
| | | BM | -173.80 | 13.75 | |
| | | psi | -173.59 | 13.96 | |
| | | EB | -171.60 | 15.96 | |
| Chaetodontidae (butterflyfishes) | PC1 | **AC** | **-194.35** | **0** | $\alpha$ = 0.19; $\sigma_{initial}$ = 3.52E$^{-07}$; root state = 0.004 |
| | | **OU** | **-194.35** | **0** | $\alpha$ = 0.10; $\sigma$ = 0.001; root state = 0.004; $t_{1/2}$ = 6.9 My |
| | | psi | -178.58 | 15.77 | |
| | | BM | -135.59 | 58.76 | |
| | | EB | -133.46 | 60.89 | |
| | PC2 | **psi** | **-269.60** | **0** | $\sigma$ = 0.0003; $\psi$ = 0.65; root state = 0.011 |
| | | AC | -263.59 | 6.01 | |
| | | OU | -263.59 | 6.01 | |
| | | BM | -242.41 | 27.19 | |
| | | EB | -240.28 | 29.32 | |
| | PC3 | **AC** | **-258.12** | **0** | $\alpha$ = 0.57; $\sigma_{initial}$ = 1.23E$^{-13}$; root state = 0.005 |
| | | **OU** | **-258.12** | **0** | $\alpha$ = 0.28; $\sigma$ = 0.002; root state = 0.005; $t_{1/2}$ = 2.5 My |
| | | psi | -246.39 | 11.73 | |
| | | BM | -208.72 | 49.41 | |
| | | EB | -206.59 | 51.54 | |
| Lutjanidae (snappers) | PC1 | **AC** | **-231.45** | **0** | $\alpha$ = 0.20; $\sigma_{initial}$ = 1.06E$^{-07}$; root state = 0.001 |
| | | **OU** | **-231.45** | **0** | $\alpha$ = 0.10; $\sigma$ = 0.002; root state = 0.001; $t_{1/2}$ = 6.9 My |
| | | BM | -206.11 | 25.34 | |
| | | psi | -205.53 | 25.93 | |
| | | EB | -204.00 | 27.45 | |
| | PC2 | **AC** | **-274.84** | **0** | $\alpha$ = 0.25; $\sigma_{initial}$ = 8.04E$^{-09}$; root state = -0.011 |
| | | OU | -269.15 | 5.68 | |
| | | psi | -256.96 | 17.87 | |
| | | BM | -239.34 | 35.49 | |
| | | EB | -237.23 | 37.61 | |
| | PC3 | **AC** | **-288.86** | **0** | $\alpha$ = 0.12; $\sigma_{initial}$ = 2.08E$^{-06}$; root state = -0.013 |
| | | **OU** | **-288.86** | **0** | $\alpha$ = 0.06; $\sigma$ = 0.001; root state = -0.013; $t_{1/2}$ = 11.6 My |
| | | psi | -281.12 | 7.75 | |
| | | BM | -278.71 | 10.16 | |
| | | EB | -276.59 | 12.27 | |
| Mullidae (goatfishes) | PC1 | **OU** | **-92.66** | **0** | $\alpha$ = 2.02; $\sigma$ = 0.049; root state = -0.007; $t_{1/2}$ = 0.3 My |
| | | AC | -86.96 | 5.70 | |
| | | psi | -74.78 | 17.89 | |
| | | BM | -38.30 | 54.37 | |
| | | EB | -36.09 | 56.58 | |
| | PC2 | **OU** | **-122.44** | **0** | $\alpha$ = 0.88; $\sigma$ = 0.014; root state = 0.008; $t_{1/2}$ = 0.8 My |
| | | **AC** | **-120.94** | **1.50** | $\alpha$ = 1.75; $\sigma_{initial}$ = 2.91E$^{-19}$; root state = 0.008 |
| | | psi | -111.99 | 10.45 | |
| | | BM | -76.40 | 46.04 | |
| | | EB | -74.19 | 48.26 | |
| | PC3 | **OU** | **-131.19** | **0** | $\alpha$ = 0.30; $\sigma$ = 0.259; root state = -0.004; $t_{1/2}$ = 2.3 My |
| | | **AC** | **-131.06** | **0.13** | $\alpha$ = 0.71; $\sigma_{initial}$ = 1.21E$^{-21}$; root state = -0.010 |
| | | **psi** | **-129.29** | **1.91** | $\sigma$ = 0.001; $\psi$ = 0.71; root state = -0.010 |
| | | BM | -89.60 | 41.59 | |

| | | | | | |
|---|---|---|---|---|---|
| | | EB | -87.39 | 43.80 | |
| Pomacanthidae (angelfishes) | PC1 | **OU** | **-144.04** | **0** | $\alpha$ = 0.38; $\sigma$ = 0.006; root state = -0.006; $t_{1/2}$ = 1.8 My |
| | | **AC** | **-144.04** | **0** | $\alpha$ = 0.76; $\sigma_{initial}$ = 4.59E$^{-17}$; root state = -0.006 |
| | | psi | -123.76 | 20.28 | |
| | | BM | -96.56 | 47.48 | |
| | | EB | -94.38 | 49.66 | |
| | PC2 | **OU** | **-164.10** | **0** | $\alpha$ = 0.65; $\sigma$ = 0.008; root state = -0.001; $t_{1/2}$ = 1.1 My |
| | | **AC** | **-163.95** | **0.14** | $\sigma_{initial}$ = 7.41E$^{-27}$; $\alpha$ = 1.30; root state = -0.001 |
| | | psi | -158.21 | 5.88 | |
| | | BM | -114.86 | 49.23 | |
| | | EB | -112.68 | 51.42 | |
| | PC3 | **OU** | **-177.60** | **0** | $\alpha$ = 0.63; $\sigma$ = 0.006; root state = 0.005; $t_{1/2}$ = 1.1 My |
| | | **AC** | **-177.39** | **0.21** | $\alpha$ = 1.26; $\sigma_{initial}$ = 2.62E$^{-26}$; root state = 0.005 |
| | | psi | -150.56 | 27.04 | |
| | | BM | -121.41 | 56.19 | |
| | | EB | -119.22 | 58.37 | |
| Pomacentridae (damselfishes) | PC1 | **OU** | **-799.53** | **0** | $\alpha$ = 1.62; $\sigma$ = 0.017; root state = -0.002; $t_{1/2}$ = 0.4 My |
| | | AC | -791.78 | 7.75 | |
| | | psi | -735.12 | 64.41 | |
| | | BM | -514.06 | 285.47 | |
| | | EB | -512.02 | 287.51 | |
| | PC2 | **AC** | **-1036.94** | **0** | $\alpha$ = 0.84; $\sigma_{initial}$ = 1.29E$^{-23}$; root state = -0.001 |
| | | **OU** | **-1036.53** | **0.41** | $\alpha$ = 1.63; $\sigma$ = 0.008; root state = -0.001; $t_{1/2}$ = 0.4 My |
| | | psi | -1001.26 | 35.67 | |
| | | BM | -799.43 | 237.51 | |
| | | EB | -797.39 | 239.55 | |
| | PC3 | **OU** | **-1097.87** | **0** | $\alpha$ = 1.45; $\sigma$ = 0.006; root state = -0.001; $t_{1/2}$ = 0.5 My |
| | | **AC** | **-1094.60** | **3.27** | $\alpha$ = 0.86; $\sigma_{initial}$ = 2.98E$^{-23}$; root state = -0.001 |
| | | psi | -1061.05 | 36.82 | |
| | | BM | -852.49 | 245.38 | |
| | | EB | -850.46 | 247.42 | |

The models are ranked from best to worst fit, according to AICc scores. ΔAIC indicates the difference in AIC scores between the candidate model and the best-fitting model. Alpha ($\alpha$) is used as a generic parameter indicator. The actual parameter is model specific. In the case of OU model, $\alpha$ is the strength of selection towards an optimum trait value. Concerning the AC model, $\alpha$ is the exponential rate change parameter. Sigma ($\sigma$) is the Brownian rate parameter. Psi ($\psi$) ranges between 0 and 1, and estimates the fraction of interspecific evolutionary divergence that is due to speciational change. For the OU model, the phylogenetic half-life ($t_{1/2}$) was calculated as $t_{1/2} = \ln(2)/ \alpha$.

**Extended Figure 1.** Illustrations of the pigmentation pattern spaces for every fish family, within the first three axes of the Principal Coordinates Analyses. Species are grouped by ecoregions illustrated by color code (A = Atlantic; CIP = Central-Indo Pacific, CP = Central Pacific, TEP = Tropical Eastern Pacific and WI = Western Indian Ocean). The 95% confidence interval ellipses are illustrated around the means.

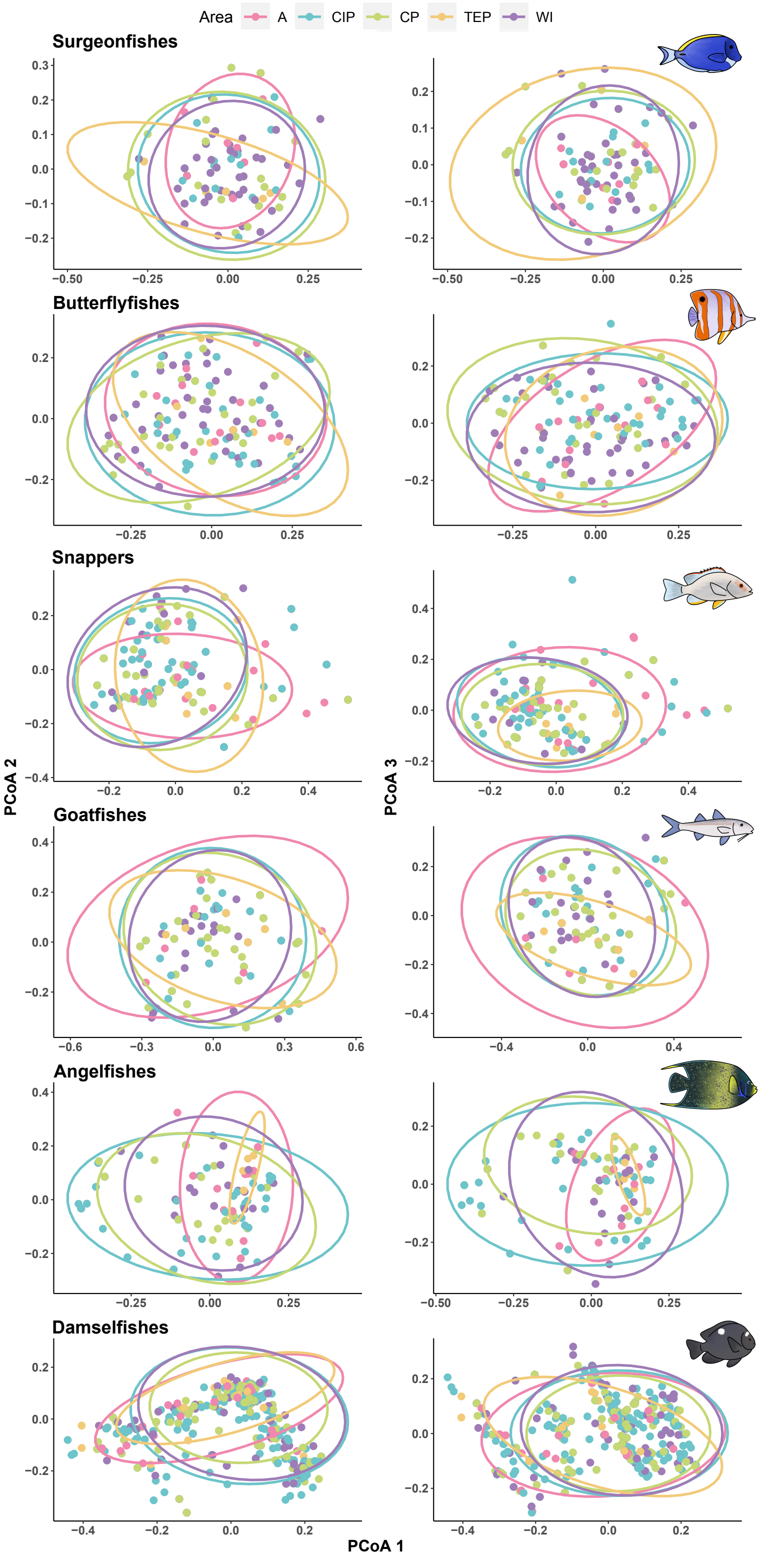

**Extended Figure 2.** Comparisons of alternative models of trait evolution using the first three axes of the Principal Coordinates Analyses summarizing the diversity of pigmentation patterns. Pie charts show the distribution of the Akaike information criterion weights (AICw) of each alternative models (BM, OU, EB, AC and psi). Values of AIC scores are provided in Extended Data Table 2. Generally, BM, EB and psi models were so poorly supported that they are not discernable on pie charts.

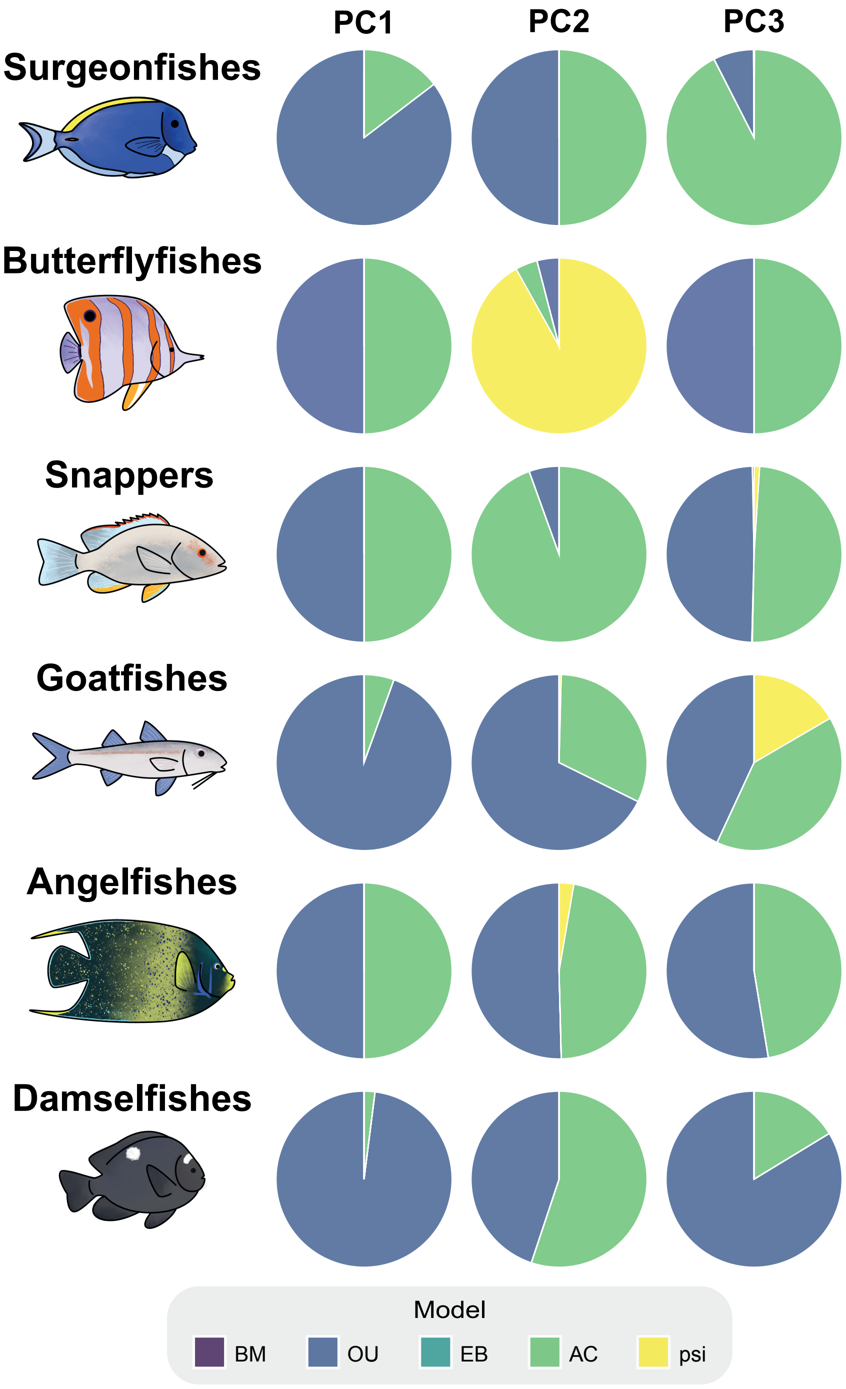

**Extended Figure 3.** Disparity through time (DTT) plots illustrating the evolution of subclade disparity across the phylogeny. Results are consistent across fish families where the subclade disparity (solid green line) was significantly higher than expected relative to the null BM model (dotted blue line with shaded area). A significant departure from BM expectation increased toward the present in all families. This demonstrates the evolution of pigmentation patterns produces greater variation within subclades than across subclades, possibly resulting in convergence.

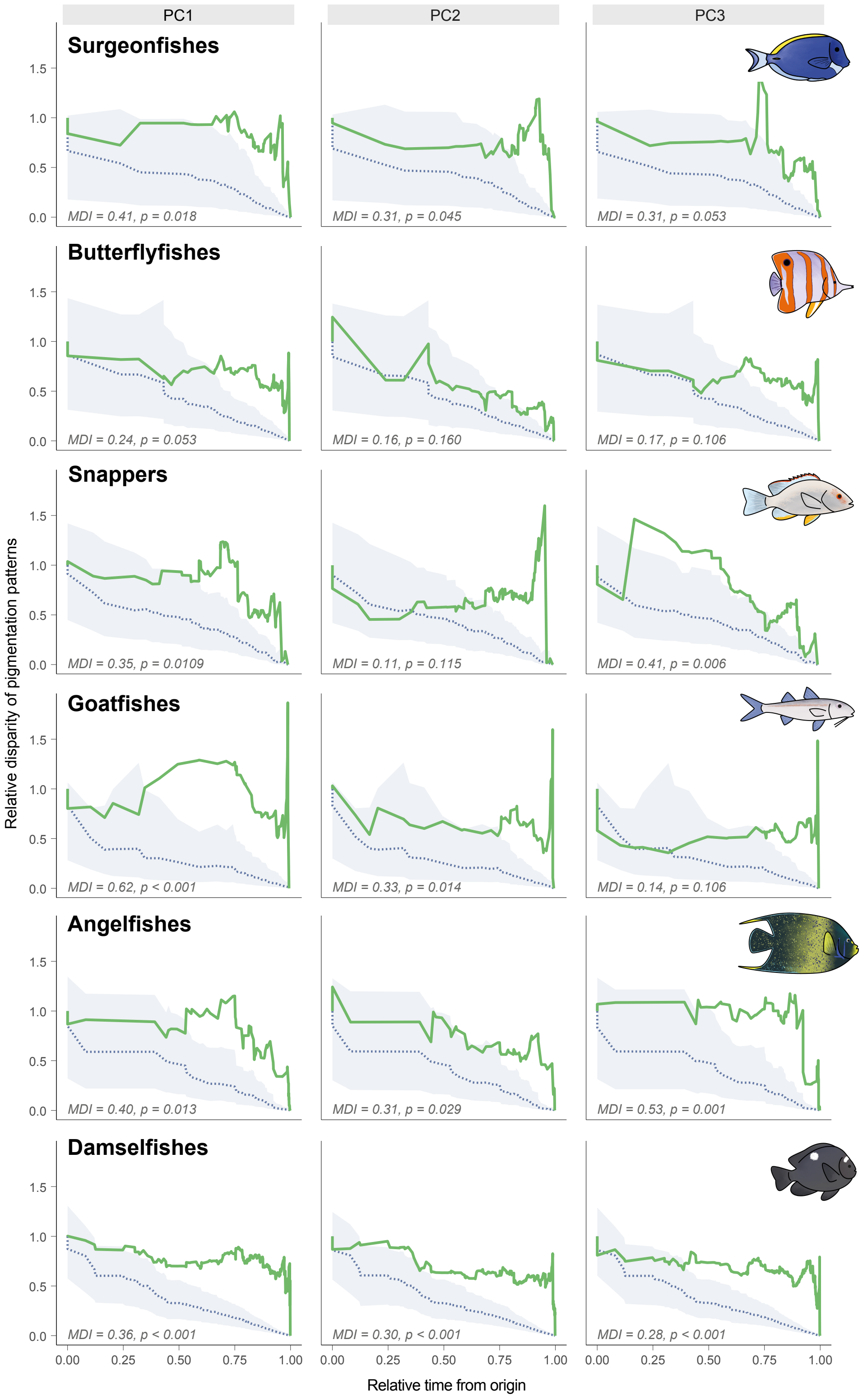

**Extended Figure 4.** Overview of the stochastic character maps (SCMs) across motif traits and fish families. Time calibrated phylogenies (top of each panel) are plotted relative to the quantiles for the times spent in each character state over the evolutionary history of each clade (bottom of each panel) across all SCMs. Character states are shaded by whether the motif was present or absent, with motif pattern organized by body region (i.e., head, trunk and tail). Quantiles are restricted to trait/branch combinations with at least 70% confidence in the SCMs. Background shading corresponds to the rise in the frequency of evolutionary transitions in motif patterns depicted in Figure 3 of the main text. This rise is comprised of numerous shifts in color pattern motifs observed below.

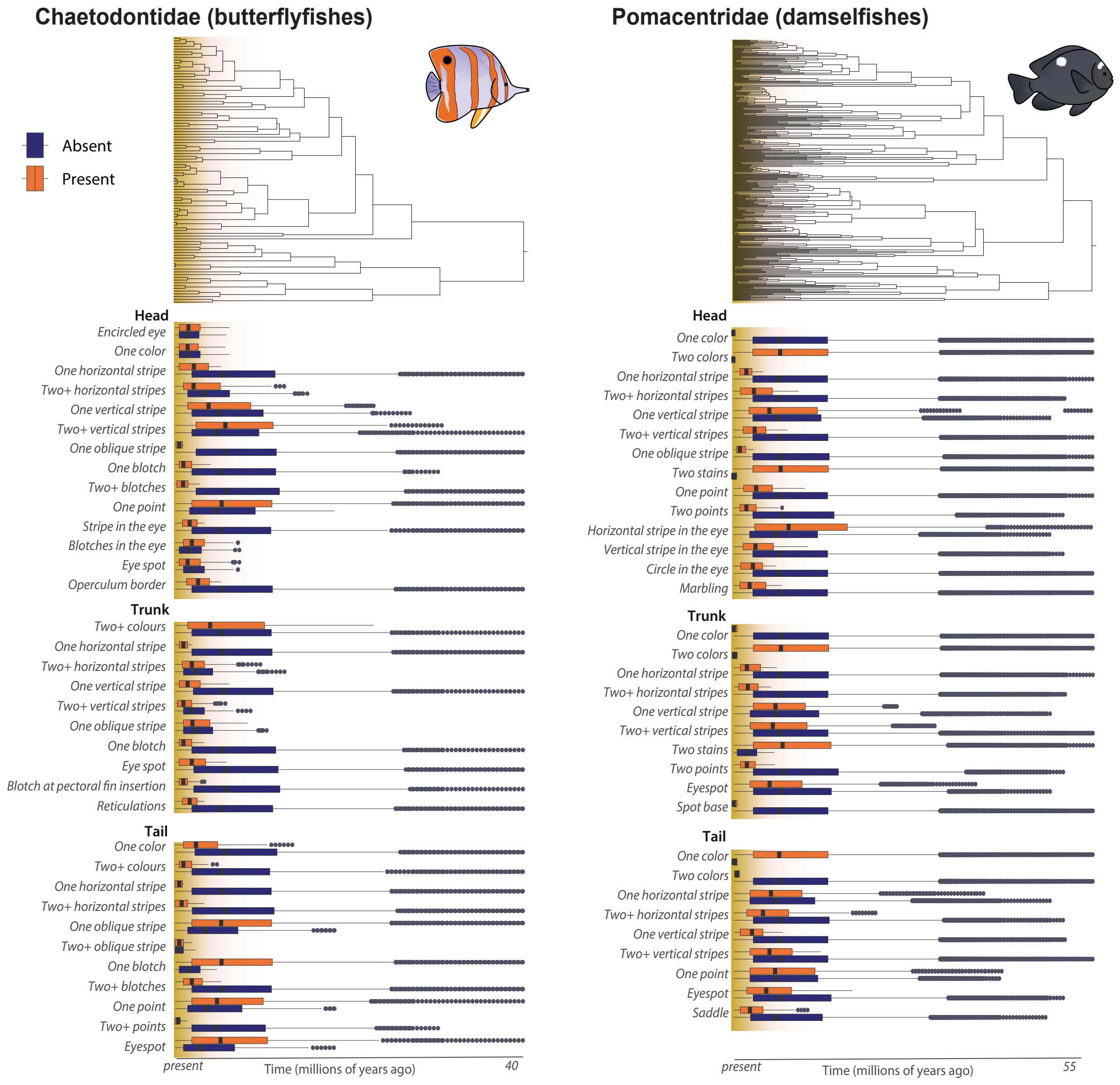

**Extended Figure 4.** (continued)

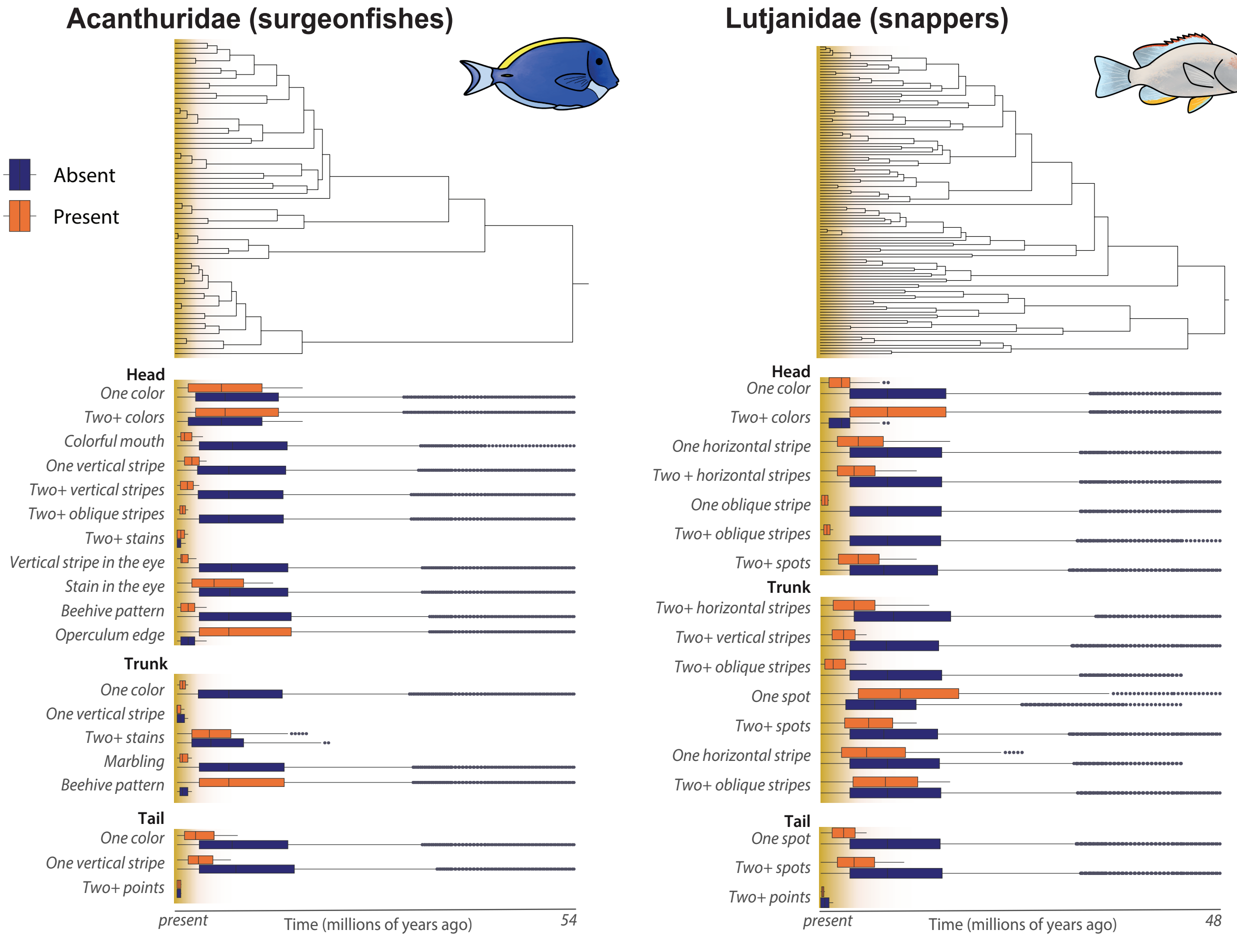

**Extended Figure 4.** (continued)

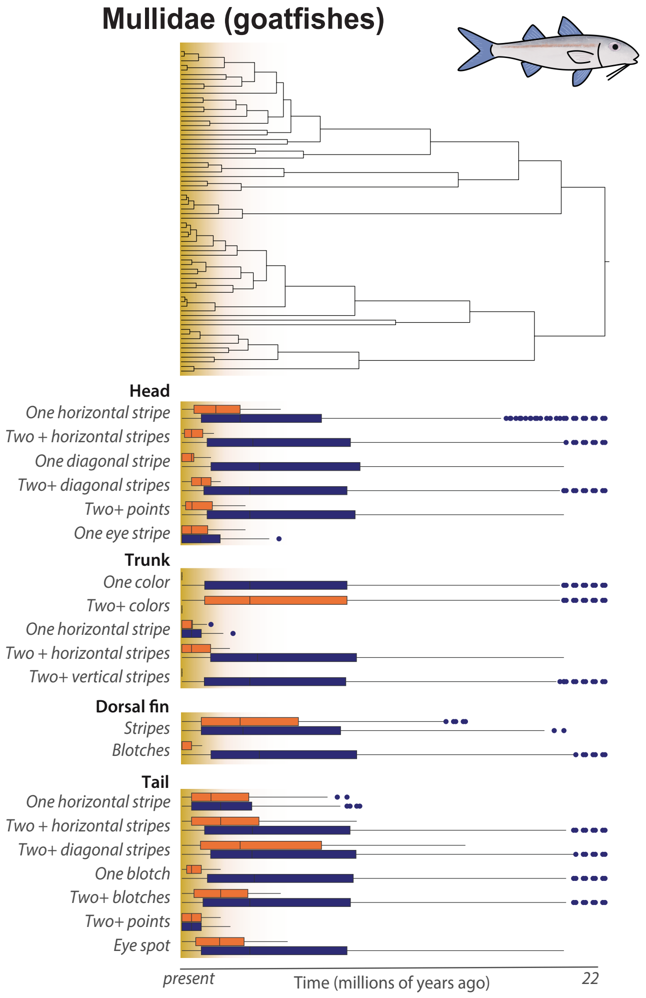


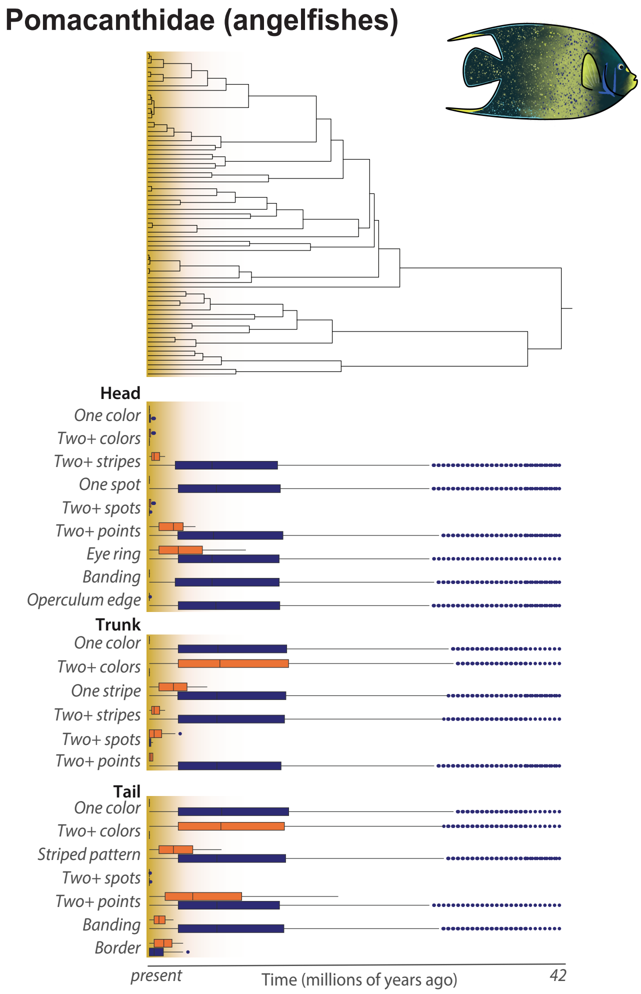
